# A CYCLIC PEPTIDE TARGETS GLIOBLASTOMA BY BINDING TO ABERRANTLY EXPOSED SNAP25

**DOI:** 10.1101/2024.09.18.613627

**Authors:** Alberto G. Arias, Laura Tovar-Martinez, Eliana Asciutto, Aman Mann, Kristina Posnograjeva, Lorena Simón Gracia, Miriam Royo, Maarja Haugas, Tambet Teesalu, Cristian Smulski, Erkki Ruoslahti, Pablo Scodeller

**Affiliations:** Medical Physics Department, Gerencia de Área Aplicaciones Nucleares a la Salud (GAANS), Centro Atómico Bariloche. San Carlos de Bariloche, Argentina; School of Science and Technology, National University of San Martin (UNSAM) and CONICET, Campus Migueletes, 25 de Mayo y Francia, CP 1650 San Martín, Buenos Aires, Argentina; Aivocode; Laboratory of Precision and Nanomedicine, Institute of Biomedicine and Translational Medicine, University of Tartu, Tartu, Estonia; Institute for Advanced Chemistry of Catalonia, IQAC-CSIC, Jordi Girona 18-26, Barcelona 08034; Biomedical Research Networking Center in Bioengineering, Biomaterials and Nanomedicine, CIBER-BBN, Jordi Girona 18-26, 08034 Barcelona; Sanford Burnham Prebys Medical Discovery Institute, 10901 N. Torrey Pines Rd, la Jolla, CA 92037, USA

## Abstract

Disease-specific changes in tumors and other diseased tissues are an important target of research because they provide clues on the pathophysiology of the disease as well as uncovering potentially useful markers for diagnosis and treatment. Here, we report a new cyclic peptide, CESPLLSEC (CES), that specifically accumulated (homed) in intracranial U87MG and the WT-GBM model of glioblastoma from intravenous (IV) injection, associating with the vasculature. Affinity chromatography of U87MG tumor extracts on insolubilized CES peptide identified Synaptosomal Associated Protein 25 (SNAP25) as a candidate target molecule (receptor) for CES. Several results supported the identification of SNAP25 as the CES receptor. IV-injected FAM-CES colocalized with SNAP25 in the tumors, and direct binding studies showed specific CES peptide binding to recombinant human SNAP25. A CES peptide-drug conjugate designed for photodynamic therapy showed selective cytotoxicity to SNAP25+ glioblastoma cell lines.

Specific accumulation of systemically injected anti-SNAP25 antibody in U87MG glioblastoma, and labeling of intact U87MG cells with anti-SNAP in flow cytometry showed that SNAP25 is available from the circulation but not in normal tissues and that it is present at the cell surface. Using an array of ECM proteins and surface plasmon resonance revealed that SNAP25 binds moderately to collagen V and strongly to collagen VI. Modeling studies suggested that CES and collagen VI compete for the same binding site on SNAP25.

Our results introduce CES as a valuable targeting peptide for drug delivery, and its receptor SNAP25 as a possible molecular marker of interest for glioblastoma.

## INTRODUCTION

Disease-specific marker molecules that are available for binding by circulating probes (“zip code” molecules)^1^ are important in two main ways. Because they are specific to a disease, they tell us something significant about that disease. Moreover, accessibility through systemic circulation makes these molecules potentially suitable as targets for diagnostic probes and drug delivery vehicles. Targeted delivery of drugs to diseased tissue has gained acceptance recently. This is particularly the case in cancer, where some antibody-drug conjugates are in the clinic and in clinical trials^2,3^. Tumor-targeting peptide-drug conjugates are also under intensive study and clinical development by academic laboratories and biotechnology companies. A tumor-penetrating peptide^4^ has produced promising phase 1 clinical trial results^5^ and is currently undergoing phase 2 controlled trials. Peptides are selective and have possibility for oral administration^6,7^.

Crucial to tumor homing is the ability of the probe to specifically recognize tumor vasculature. Tumors are not the only pathological conditions that express specific markers; other diseases also put specific molecular signatures on the vasculature of the affected tissues. The vasculature and extracellular space of diseased tissue can be probed *in situ* using *in vivo* screening of peptide libraries expressed on phage. This method has been successfully used to identify homing peptides for tumors and other diseases^8,9^, and even peptides that specifically home to different organs or tissues^10^.

We previously used *in vivo* phage display to search for peptides recognizing Alzheimer’s Disease (AD) brain^9^. One such peptide, cyclic CDAGRKQKC has been reported and was shown to recognize the vasculature of AD brain very early in the development of the disease. The target molecule for the peptide is connective tissue growth factor (CTGF) bound to the vascular extracellular matrix^9^. In studying another peptide from this the same phage screen, we noticed that this previously unreported peptide not only recognized AD brain vasculature but strongly accumulated in glioblastoma tumors, associating primarily with the blood vessels. We decided to characterize the molecular change defined by this peptide by studying tumor tissue, which are more readily available than AD brain tissue. Here, we identify the target molecule (receptor) for the peptide and report on some of its properties.

## RESULTS

### An AD-peptide homes to glioblastoma tumors

An *in vivo* phage library screen in a genetically modified mouse AD model^9^ identified a cyclic 9-amino acid peptide CESPLLSEC (CES) as a peptide that specifically accumulates (homes) in the blood vessels of AD brain (will be reported elsewhere). Analysis of the disease specificity of the CES peptide showed that FAM-labeled CES also homed to U87MG tumor tissue after 1 hour of administration (Fig. 1A) co-distributing with the blood vessel marker CD31 (insets in Fig. 1A). We also evaluated FAM-CES in the WT-GBM xenograft model. This model was developed by Blouw *et al*. ^11,12^ by isolating astrocytes from the hippocampus of neonatal mice and transforming them with the oncogenes SV40 large T-antigen and V12–H-ras. FAM-CES also targeted WT-GBM tumors from intravenous administration (Fig. 1B) and also showed colocalization with CD31 (inset in Fig. 1B).

**Figure 1.**
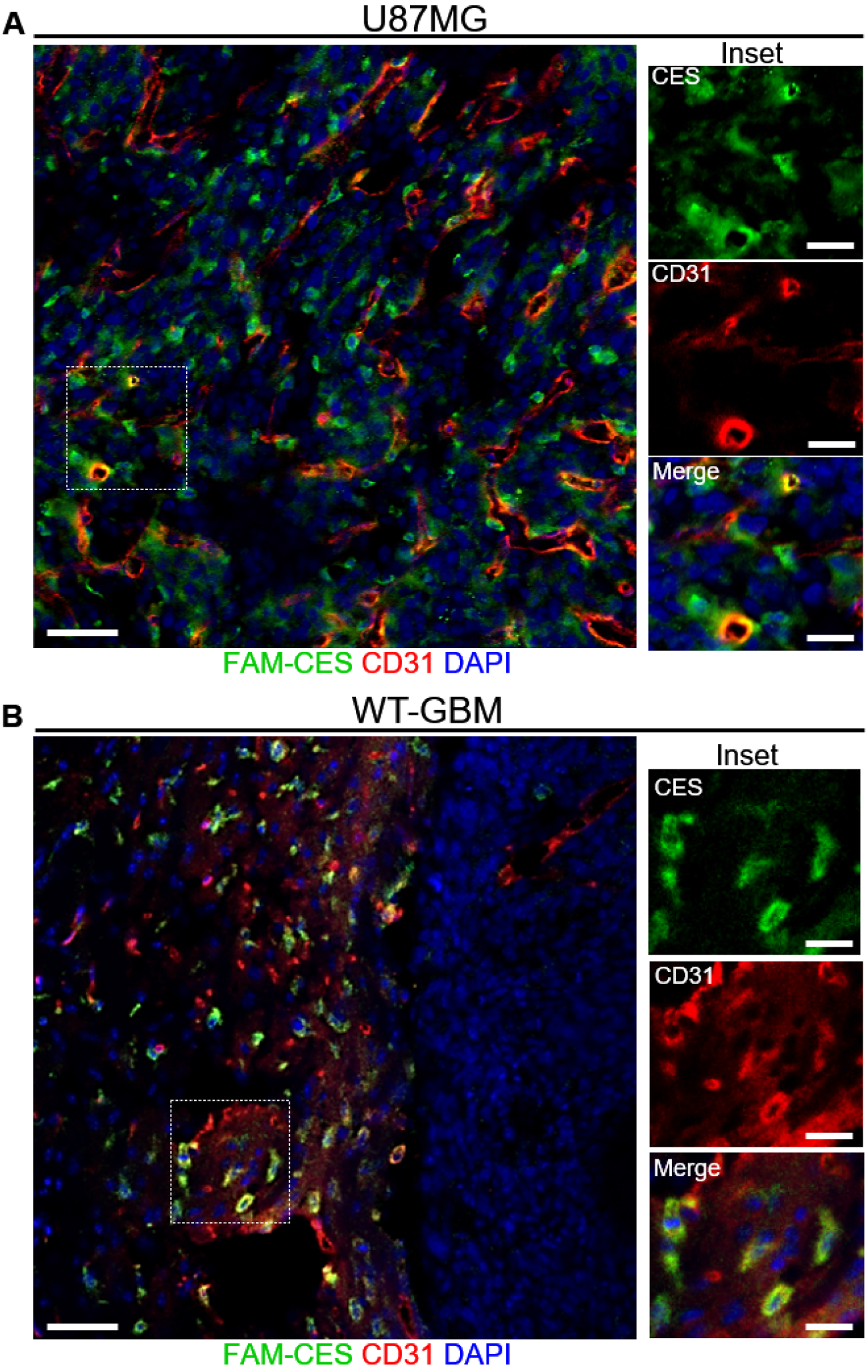
FAM-CES homes to intracranial U87MG and WT-GBM glioblastoma and colocalizes with blood vessels. Homing of intravenously administered FAM-CES in mice bearing intracranial U87MG **(A)**, WT-GBM **(B)** or orthotopic MCF10CA1a tumors **(C)** (n=3). The insets in A and B show colocalization between FAM-CES and CD31 in the tumor. 30 nmoles of FAM-CES were administered intravenously and allowed to circulate for 60 min. The mice were perfused, and the brains were fixed, sectioned, and stained for FAM (green) and CD31. Scale bars represent 50µm, scale bars for the insets represent 20µm.

### Affinity chromatography identifies the SNARE protein SNAP25 as a CES receptor

To identify a receptor of CES, we used affinity chromatography on streptavidin-coated magnetic beads functionalized with biotin-CES. U87MG tissue, which exhibited robust binding of FAM-CES in the homing studies, was used as starting material. Mass spectrometry analysis of the fractions recovered from the affinity matrix with an excess of free CES peptide, revealed several hits (Table 1 in supplementary information). We excluded cytosolic proteins and focused on synaptosomal-associated protein 25 (SNAP25) as a possible receptor because it had a high CES-eluted/control peptide-eluted ratio, and because it is overrepresented at the RNA level in the U87MG cells (Human Protein Atlas). SNAP25 is a member of the SNARE (soluble N-ethylmaleimide-sensitive factor attachment protein receptor) complex which regulates synaptic vesicle formation. According to the protein atlas, SNAP25 is expressed preferentially in the central nervous system. At the RNA level, it is expressed mainly in the brain and to a much lesser extent in eye and endocrine tissue. In cancer, according to the curated cancer atlas^13^, SNAP25 RNA is expressed in brain tumors and, outside the CNS, in neuroendocrine tumors. In line with this, we observed no tumor homing of FAM-CES in a model of orthotopic breast cancer (Fig. S1).

We first analyzed SNAP25 expression and its colocalization with intravenously injected FAM-CES, in the U87MG. FAM-CES showed apparent colocalization with SNAP25 in U87MG tumors (Fig. 2A) and no homing to healthy brain (Fig. 2B).

**Figure 2.**
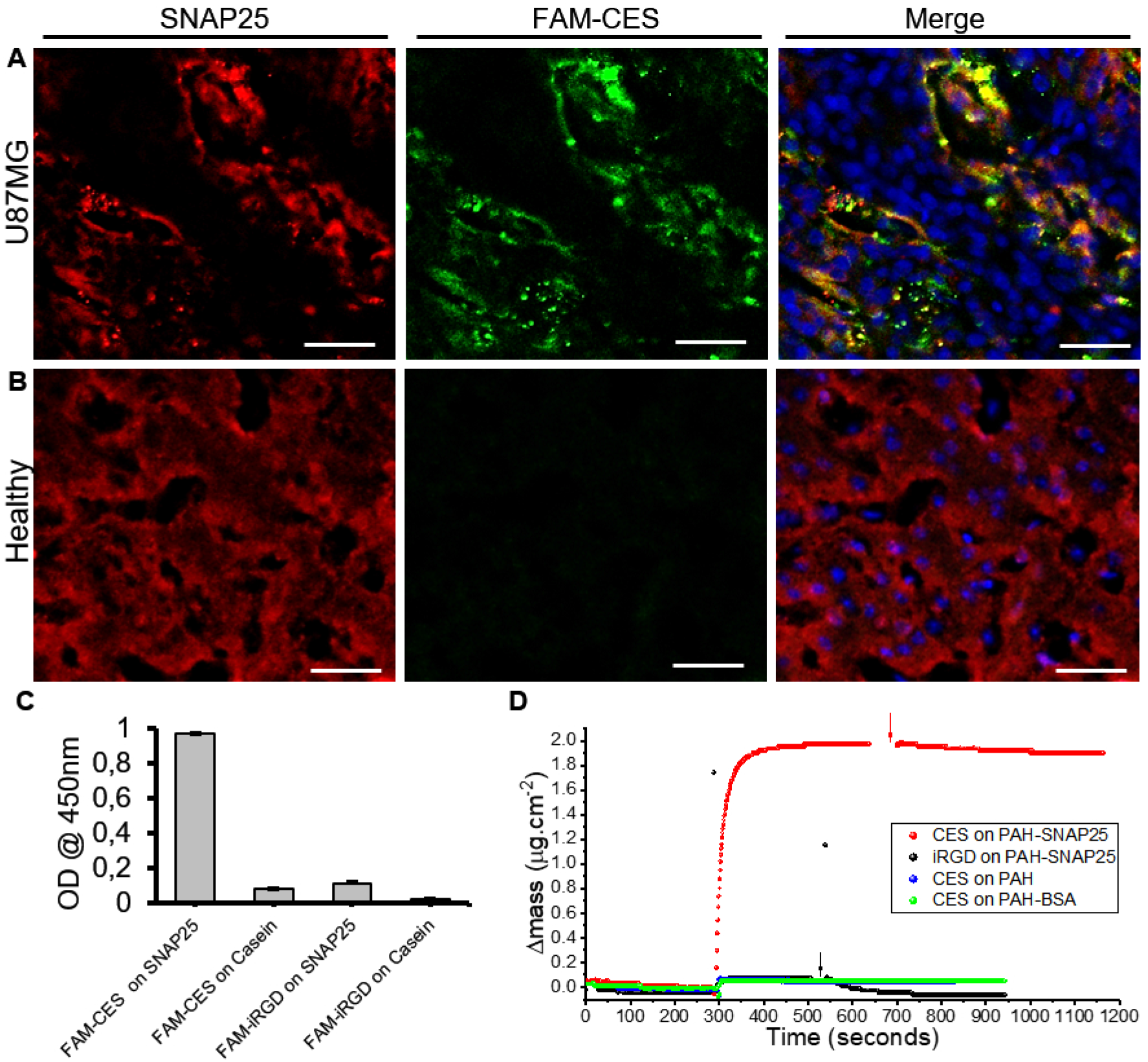
CES targets and binds to SNAP25. Homing of intravenously administered FAM-CES in mice bearing intracranial U87MG **(A)**, or in healthy mice **(B)**. 30 nmoles of FAM-CES were administered intravenously and allowed to circulate for 60 min (n=3). The mice were perfused, and the brains were fixed, sectioned, and stained for FAM (green) and SNAP25 (red). **(C)**: ELISA binding assay of FAM-CES and FAM-iRGD on plates coated with SNAP25 or Casein. **(D):** QCM analysis of the mass changes brought about by CES binding to human recombinant SNAP25 and subsequent dissociation. SNAP25 was immobilized on a viscoelastic film^20^ deposited on a gold-modified quartz crystal resonating at 5 MHz. The deposition of the protein was accomplished by using layer-by-layer self-assembly on a layer of polyallylamine (PAH)^21 22 23^. The figure shows association and dissociation of CES (red) or iRGD (black). The final concentration of peptides was 30µM. CES did not bind to PAH (blue) or to the PAH-BSA multilayer (green). Washes with PBS are denoted by arrows. Scale bars=50µm.

We also observed that SNAP25 was unevenly in U87MG brain; as opposed to the homogeneous distribution in healthy brain (Fig. 2B, red). Subnormal expression levels of SNAP25 have been reported in glioblastoma^14,15^.

We subsequently analyzed binding of FAM-CES to recombinant SNAP25, using ELISA. This experiment showed that FAM-CES bound to SNAP25 but not to Casein, and that another peptide with similar molecular weight and isoelectric point, FAM-iRGD, did not bind to SNAP25 (Fig. 2C). To evaluate if label-free CES also binds to SNAP25, and to estimate an affinity constant, we then analyzed the binding of CES to recombinant SNAP25 using Quartz Crystal Microbalance (QCM), a powerful technique to study label-free ligand-receptor interactions in solution^16,17,18,19^. Our experiments showed that CES, but not iRGD, bound to SNAP25, and that CES did not bind to the control layers poly(allylamine hydrochloride) (PAH) or multilayer PAH-BSA (Fig. 2D). CES, showed a slow dissociation rate upon washing with PBS (Fig. 2D, black arrow). Association and dissociation fitted well to a one-to-one model and yielded an estimative K_D_ of 24nM.

### A CES-Verteporfin conjugate (CES-V) is selectively cytotoxic to SNAP25^+^glioblastoma cells

According to the protein atlas, the expression of SNAP25 in tissues is specific to the brain and comes from neuronal and glial cells. Hence, here we first evaluated expression of SNAP25 in glioblastoma and neuroblastoma cell lines. Western blot revealed that the glioblastoma cell lines U87MG and LN229, but not the neuroblastoma cell line SK-N-AS, express SNAP25 (Fig. 3A)

**Figure 3.**
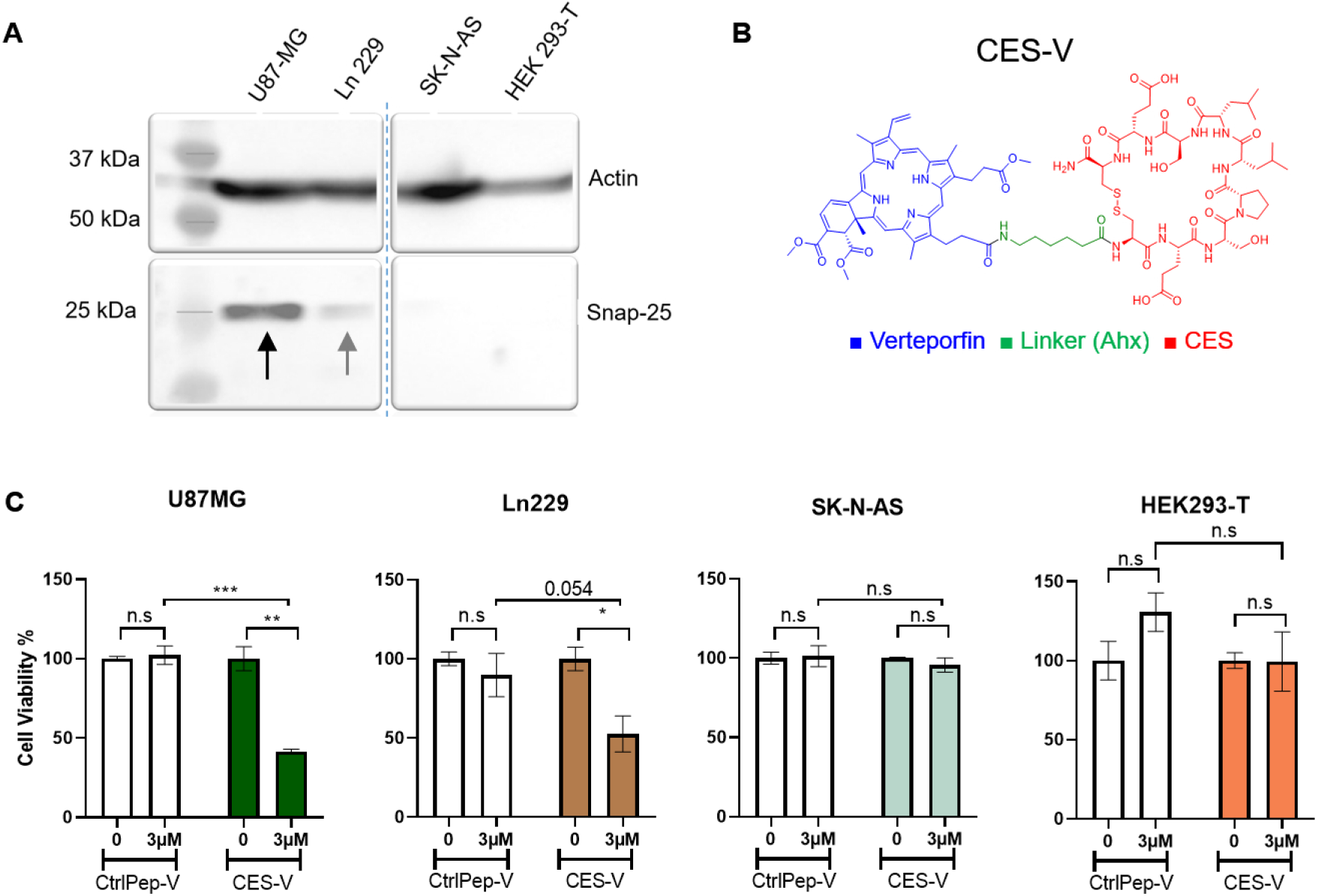
Cytotoxicity of irradiated CES-Verteporfin conjugate (CES-V) to cultured brain cancer cells. **(A):** Western blot showing expression of SNAP25 in the different cell lines. **(B):** Structure of the peptide-drug conjugate CES-V, with the peptide part in red, the linker in green and the drug in blue. **(C):** Cultured U87MG, LN229, SK-N-AS, or HEK293-T cells were incubated with 3μM of CES-V or CtrlPep-V for 30 min at 37°C, followed by three washes with media. Then 100µL of new media was placed, and the wells were irradiated with a dose of 10 J/cm^2^ (using an irradiance of 160 mW/cm^2^) using a 690nm laser. After irradiation, the cells were cultured for an additional 24 h, and the viability was assessed by MTS assay.

To see if CES would have the potential to deliver drugs to SNAP25^+^ cells, we designed a CES-drug conjugate (Fig. 3B) and tested the conjugate for cytotoxicity to cultured brain cancer cells from different cell lines. For this, we sought to use a photosensitizer, which eliminates the need for a cleavable linker. As a photosensitizer, we picked Verteporfin, as it contains a carboxylic acid for easy conjugation to the N-terminus of the peptide, is FDA-approved, and absorbs in the far-red (690nm), a spectral region with low absorption and scattering in tissues. We incubated the cells with the CES-Verteporfin (CES-V) or Control peptide-Verteporfin (CtrlPep-V) conjugate and evaluated cell viability with and without irradiation. Irradiated CES-V showed significant toxicity in U87MG cells and LN229 cells and showed no toxicity to SK-N-AS or HEK293T cells (Fig. 3C). The difference in toxicity between irradiated CES-V and irradiated CtrlPep-V only reached significance for U87MG, correlating with the SNAP25 expression levels of those cells. The CES peptide alone, even at concentrations as high as 30µM did not elicit cytotoxicity in the U87MG cells (Fig. S2), indicating that Verteporfin and irradiation are required for the cytotoxicity.

### SNAP25 becomes accessible *in vivo* in U87MG tumors

For SNAP25 to be the CES receptor *in vivo*, it should be accessible to ligands from the bloodstream. To study SNAP25 accessibility *in vivo*, we injected anti-SNAP25 antibody to U87MG tumor-bearing and healthy control mice. The antibody accumulated in the tumor, but not in the contralateral side of the brain, or in the brain of healthy mice (Fig. 4A). Parallel staining detected only marginal amount of endogenous mouse IgG (concentration in mouse blood: 5-9 mg/mL^24^), in glioblastoma (Fig. 4B left panel), indicating that the anti-SNAP25 homing was specific and not a consequence of tumor leakiness. Co-staining for CD31 and administered anti-SNAP25 showed that the antibody partially targeted the blood vessels (Fig. 4B, right panel, white arrows). The tumors of noninjected U87MG mice showed punctate SNAP25 staining on their blood vessels (Fig. 4C and Fig. S3).

**Figure 4.**
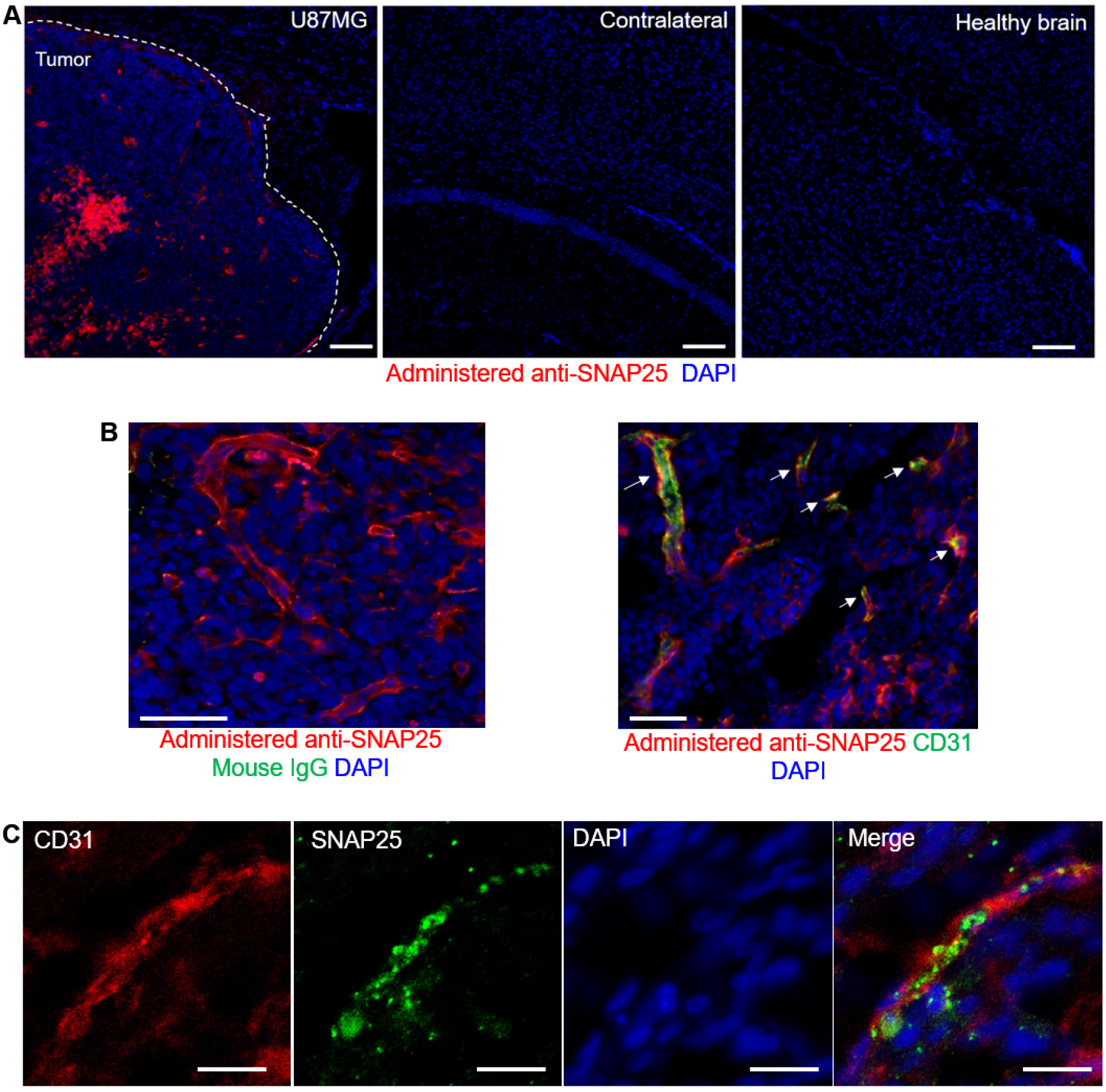
SNAP25 in U87MG tumors is accessible to circulating antibody. **(A):** Rabbit anti-SNAP25 was administered to mice bearing intracranial U87MG tumors and allowed to circulate for 24 hours. The mice were then perfused, and their tissues fixed, sectioned, and stained for rabbit IgG. **(B):** Administered anti-SNAP25 (red) together with endogenous mouse IgG (green, left panel) or CD31 (green, right panel) in the tumor. **(C):** SNAP25 and CD31 costaining in U87MG tumor of un-injected mice. Scale bars for A and B represent 100µm, scale bars in C represent 20µm. Representative images from n=3 mice are shown.

### SNAP25 is atypically exposed in U87MG and binds to collagens V and VI

In non-pathological conditions, SNAP25 is on the inner leaflet of the plasma membrane. Since SNAP25 appeared to be accessible to FAM-CES *in vivo* and to CES-V *in vitro* in U87MG, we hypothesized that it would perhaps be exposed on the outer leaflet of the plasma membrane. To test this, we performed live cell flow cytometry of U87MG cells at 4°C, in the absence of any detergent, to evaluate superficial SNAP25. Our results showed a SNAP25^+^ population (approximately 50%) in the U87MG cells, that was not seen in the control cells (Fig. 5A). We also detected SNAP25 in confocal microscopy, on cultured, fixed, U87MG cells, stained without any detergent (Fig. 5B). Of note, evaluation of cell-surface SNAP25 expression in a cell line of neuroendocrine cancer (the other SNAP25-expressing cancer in humans) also showed a SNAP25^+^ population (Fig. S4).

**Figure 5.**
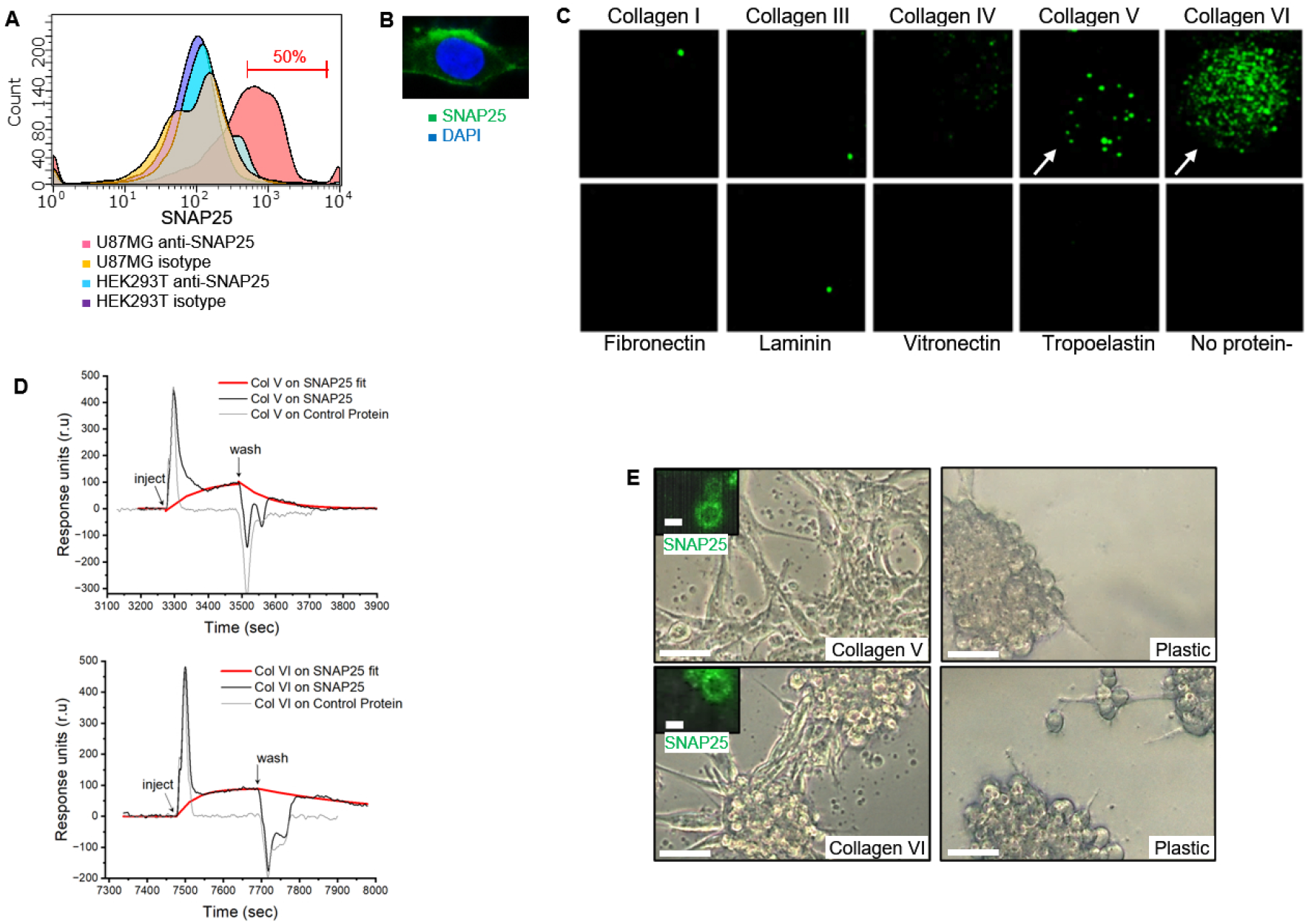
SNAP25 is exposed on the cell surface and binds to collagens V and VI. **(A):** Live cell flow cytometry without permeabilization to detect superficial SNAP25. Cultured U87MG or HEK293Tcells were stained for SNAP25 using A488 anti-SNAP25 or its isotype control. **(B):** Immunofluorescent confocal microscopy of cultured U87MG cells, stained in the absence of any detergent (green: SNAP25, blue: DAPI). **(C):** Binding of SNAP25 to ECM proteins. Recombinant SNAP25 was added to an ECM array and A488 anti-SNAP25 was used to detect binding. **(D):** Surface plasmon resonance sensorgram of collagens V and VI on sensors coated either with recombinant SNAP25 or a control protein of similar molecular weight. **(E):** Optical microscopy shows **c**ellular morphology on collagen V and VI of cells seeded on either collagen V/VI coated plates or uncoated plates (scale bars= 20μm); the inset shows SNAP25 immunofluorescence of the vesicle-resembling particles (scale bars= 2μm).

Because another SNARE protein, Syntaxin-4, has been found to translocate to the surface in teracarcinoma cells^25^ and to bind the extracellular matrix proteins laminin and syndecan-1^26^, we hypothesized that SNAP25 could also interact with ECM proteins. To test this, we evaluated the binding of recombinant SNAP25 to an array of different ECM proteins (collagen I, III, IV, V, VI, fibronectin, laminin, vitronectin, tropoelastin). This experiment showed binding of SNAP25 to collagens V and VI (Fig. 5C, white arrows). We then evaluated binding of collagens V and VI to recombinant SNAP25 immobilized via His-Tag/Ni-NTA chemistry, using surface plasmon resonance. This experiment confirmed interaction of collagens V and VI to SNAP25 and no binding to a control protein of similar molecular weight and immobilized in the same way (Fig. 5D). The data fitted well to a one-to-one interaction model yielding estimative K_D_ binding constants of 7.2µM and 15nM for collagen V and VI respectively.

Given this finding, we hypothesized that a SNAP25-Collagen V/VI interaction could be participating in cell attachment. U87MG cells grown on collagens V and VI formed monolayers and presented particles attached to the plate, as opposed to the same cells grown on plastic, which formed colonies and did not display these particles attached to the plate (Fig. 5E). Closer analysis of these particles using confocal microscopy and SNAP25 immunostaining revealed a vesicle-resembling structure of approximately 3μm and positivity for SNAP25 (inset in Fig. 5E).

### Modeling suggests that CES and Collagen VI compete for SNAP25

We wondered if CES could bind to the same region on SNAP25 as collagen. To investigate this possibility, we performed computational studies to obtain an idea on the binding sites for CES and collagen VI on SNAP25.

SNAP25 is composed of three main regions or domains: two SNARE motifs, with alpha helix structures, in the regions 1-82 (“SN1”, blue in Fig. 6A left), and 142-206 (“SN2”, red in Fig. 6A left), and a third region between 83-141, an unstructured helix referred to as the linker region (yellow/cyan in Fig. 6A left). The SNARE motifs interact with other SNARE proteins to form the synaptic vesicle. The linker region contains a palmitoylation site, composed of 4 cysteines (Cys84, 85, 90, 92)^27^; palmitoylation is needed for SNAP25 to bind to the plasma membrane^28^. Moreover, it has been shown that the motif 116-120 of the linker region, QPARV, facilitates palmitoylation^28^.

**Figure 6.**
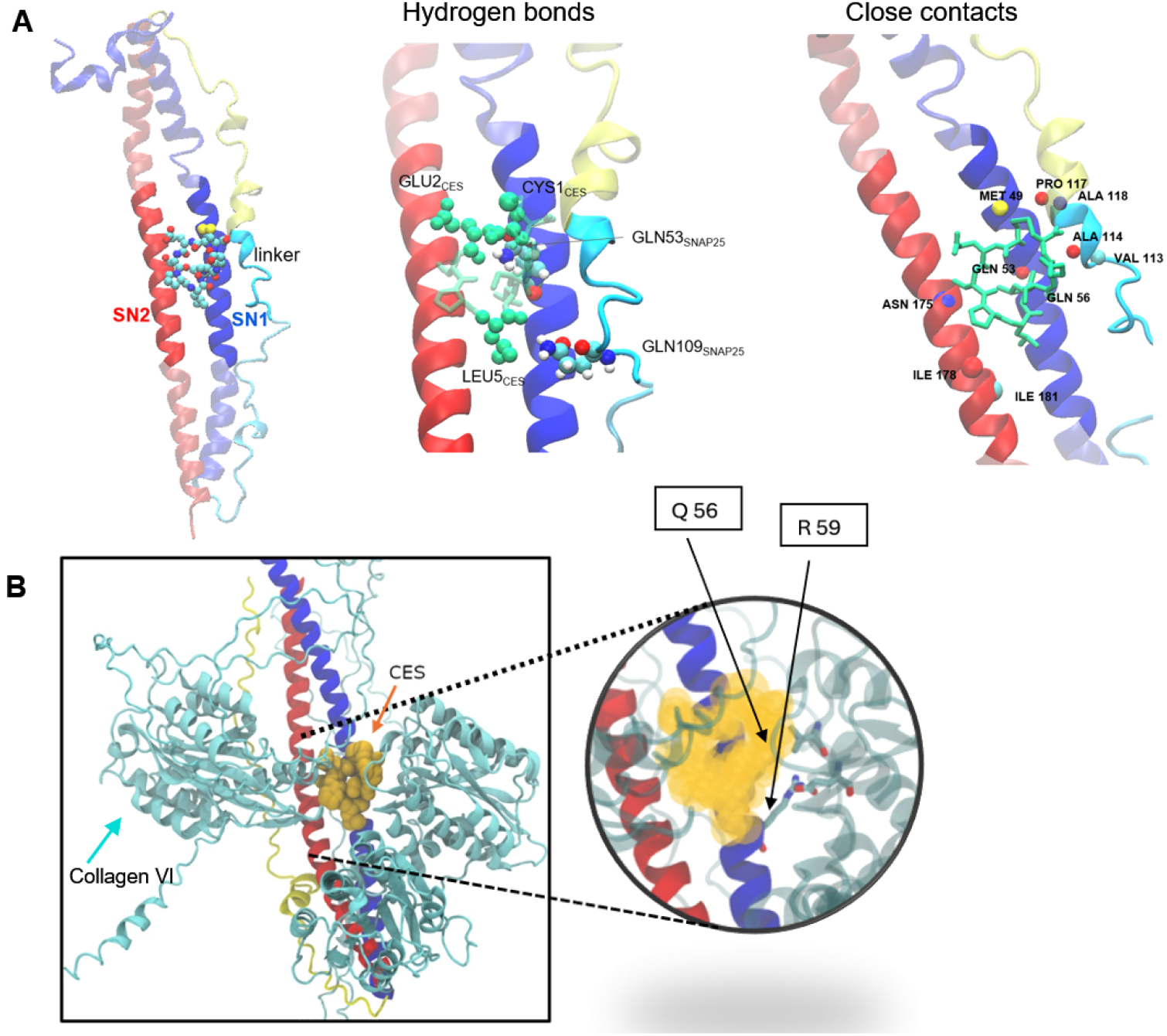
Modeling of SNAP25-CES and -collagen VI complexes. **(A):** Representation of SNAP25 and the best CES pose obtained by docking (left panel). Hydrogen bonds formed between GLN53_SNAP25_ - GLU2_CES_, GLN53_SNAP25_ - CYS1_CES_, and GLN109_SNAP25_ - LEU5_CES_ (middle panel), and close contacts between SNAP-25 and CES in sphere representation (right panel). **(B)**: SNAP25-collagen VI complex with CES superimposed in yellow spheres highlights competition for the same binding site. A close-up to the binding site reveals interactions between collagen VI and SNAP25, specifically involving residues Q56 and R59.

Docking showed that the CES pose with the best score was in a region between SN1 and SN2, making close contacts with both helices and a portion of the linker (Fig 6A, left panel). This pose was subjected to molecular dynamics for refinement and to obtain more details on the physical interactions governing the binding process. We found that on average, CES bound to SNAP25 forming three hydrogen bonds. Residue GLN53_SNAP25_ from SN1 formed H-bonds with GLU2_CES_ and CYS1_CES_, while a third H-bond is formed between GLN109_SNAP25_ and LEU5_CES_ and connects the peptide to the linker (Fig. 6A, middle panel). The two H-bonds formed with SN1 (GLN53_SNAP25_) were present 89 % of the simulation time, whereas the H-bond with the linker (GLN109_SNAP25_) was less frequent (16 % of the simulation time). Close contacts were found along the three domains (residues 49-56, 113-118, and 175-181), with the shortest distances being to residues 1-3 of CES (Figure 6A, right panel).

We then modelled the interaction of SNAP25 with the strongest binding collagen (VI) and compared it with the CES docking pose. This showed that CES and Collagen VI occupy the same region, specifically between SN1 and SN2, and that collagen VI closely interacts with residues Q56 and R59 (Fig. 6B), residues that are in the CES docking pose; suggesting that both collagen VI and CES compete for the same binding site on SNAP25. In line with this, free CES affected the adhesion of U87MG cells to collagen VI-coated plates (Fig. S5).

The binding motif within collagen VI contains the DXXYGE motif, which is present in the SNAP25-binding collagens (V and VI) and absent in the non-binding ones (I, III, IV).

## DISCUSSION

An attractive aspect of testing different candidate targeting peptides from one same phage screen is that the peptides often recognize separate receptors, leading to discovery of different circulation-accessible biomarkers. Here, we found that CES homes to glioblastoma in mice. We observed this phenomenon before with the other targeting peptide from this screen (DAG) which showed targeting to a patient derived model of glioblastoma^9^.

Affinity chromatography identified SNAP25 as a CES receptor. SNAP25 is a member of the SNARE complex, which regulates synaptic vesicle assembly and neurotransmission. In addition to SNAP25 emerging from the CES affinity chromatography as the most likely receptor candidate, tissue colocalization of fluorescence from FAM-CES with SNAP25 immunostaining, and direct binding of FAM-CES and CES to recombinant SNAP25 (by ELISA, and quartz crystal microgravimetry, respectively), showed that SNAP25 is a CES receptor.

To explore if SNAP25 can be exploited for drug targeting we designed a simple, proof-of-concept, peptide-drug conjugate, by coupling CES, to the FDA approved photosensitizer Verteporfin. Irradiated CES-V showed cytotoxicity to the SNAP25^+^ glioblastoma cell lines tested but not to the SNAP25^-^ neuroblastoma one.

We inferred that the ability of SNAP25 to act as a CES receptor likely requires extracellular exposure to be available to ligands such as CES or anti-SNAP25 antibodies. Indeed, flow cytometry of U87MG cells revealed extracellular exposure of SNAP25 and, in vivo, SNAP25 became accessible to a circulating antibody only in glioblastoma tissue.

We observed SNAP25^+^ tumor blood vessels. In endothelial cells (according to the protein atlas) SNAP25 RNA is not detected, and at the protein level, SNAP25 could not be detected in cultured endothelial cells^29^. In cancer (according to the curated cancer atlas), low levels of SNAP25 RNA are detected in endothelial cells of brain tumors and neuroendocrine tumors; and none in fibroblasts. Based on this, it is possible that the vascular SNAP25 we observed comes from adhered SNAP25^+^ particles secreted by U87MG cells.

Aberrant expression of intracellular proteins at the cell surface in proliferating or otherwise activated cells, such as cancer cells, is a well-documented, albeit poorly understood phenomenon. These are ubiquitously expressed intracellular proteins in normal resting cells but are present and available for ligand binding in activated cells. Examples include: nucleolin in tumor vessels^30^, gC1qR/p32 in tumor endothelial cells, tumor cells, and tumor-associated and other macrophages^31,32^. Among SNARE proteins, Syntaxin-4 has been found to translocate to the surface in teracarcinoma cells^25^. Cell surface SNAP25 has, to our knowledge, not been reported before; however, SNAP25 has been detected previously by other researchers on the surface of extracellular vesicles secreted by activated astrocytes^33^. Thus, SNAP25 appears to belong to the general group of proteins that are aberrantly exposed in disease. Some functional activities have been assigned to the aberrantly exposed intracellular proteins^34^.

Another relevant discovery of this work is the interaction found between SNAP25 and collagens V and VI, which, to our knowledge, has not been described before. Collagen VI is an accessible biomarker in U87MG tumors^35^ and is expressed on pathological blood vessels in human glioma^36^, and glioblastoma cells secrete collagen VI to facilitate invasion^37^. Whether blocking SNAP25 or disrupting the SNAP25-collagen VI interaction using CES or an antibody will have a therapeutic effect in glioblastoma will be subject of a further study.

Our results introduce a useful targeting peptide for glioblastoma, CES, which can be used for drug delivery, and SNAP25 as a possible new drug delivery target in glioblastoma.

## Acknowledgements

This project was started with support from the US Defense Advanced Research Projects Agency (DARPA) under Cooperative Agreement HR0011-13-2-0017. The content of the information within this document does not necessarily reflect the position or the policy of the US Government. ER and by grant R21 AG058013 from the US National Institute of Aging. P.S. acknowledges support from MICIN/AEIMCIN/AEI/10.13039/501100011033/ and by “ERDF A way of making Europe”, (grant PID2021-122364OA-I00; and Ramon y Cajal contract RYC2020-028754-I), the Estonian Research Council (grant PUT PSG38). A.G.A acknowledges a PhD fellowship from CNEA (Argentina) and a Dora Plus doctoral visiting fellowship from the University of Tartu. L.T-M. acknowledges a JAE Intro fellowship from CSIC. M.R. acknowledges support from CIBER BBN(CB06/01/0074) and by the Generalitat de Catalunya (2021SGR00230). CIBERBBN is financed by the Instituto de Salud Carlos III with assistance from the ERDF. We also acknowledge the ICTS NANBIOSIS for the support of the Synthesis of Peptides Unit (U3) at IQAC-CSIC and Gerardo Acosta for peptide synthesis. TT was funded by the Estonian Research Council (grants PRG230 and PRG1788), EuronanomedIII projects ECM-CART and iNanoGun, and TRANSCAN3 project ReachGLIO. C.S. was funded by the Agencia Mincyt (Argentina) (projects PICT-2021-I-A-00865)

## MATERIALS AND METHODS

### Cells

U87MG cells (ATCC HTB14) LN-229 (ATCC CRL-2611), HEK293T (CRL-3216), SK-N-AS (ATCC CRL-2137) were grown in DMEM F12 (Lonza, Belgium) containing 100 IU/mL of penicillin and streptomycin (Gibco), Amphotericin B (0.25 µg/ml), 10% of heat-inactivated fetal bovine serum (Gibco, USA or NATOCOR, Argentina). Cells were cultured at +37 °C in a humidified atmosphere containing 5% CO_2_. MCF10CA1a human triple-negative breast cancer cells (obtained from the Erkki Ruoslahti laboratory at the Cancer Research Center, Sanford Burnham Prebys Medical Discovery Institute) were cultured in DMEM supplemented with 100 IU/mL streptomycin, penicillin, and 10% FBS.

### Peptides and peptide-drug conjugates

Unlabeled peptides or FAM or biotin labeled peptides were purchased from TAG Copenhagen.

| Notation used | Peptide | Notes |
| --- | --- | --- |
| CES | Ac-CESPLLSEC-NH<sub>2</sub> | Disulfide: C1-C9 |
| iRGD | Ac-CRGDKGPDC-NH<sub>2</sub> | Disulfide: C1-C9 |
| FAM-CES | FAM-Ahx-CESPLLSEC-NH<sub>2</sub> | Disulfide: C1-C9 |
| FAM-CTRL | FAM-Ahx-CASPRLSKC-NH<sub>2</sub> | Disulfide: C1-C9 |
| CES-V | Vert-Ahx-CESPLLSEC-NH<sub>2</sub> | Disulfide: C1-C9 |
| Ctrl-V | Vert-Ahx-AKPCGS-OH |  |

CES-V was prepared in the Peptide Synthesis Unit (U3) at IQAC-CSIC (https://www.nanbiosis.es/portfolio/u3-synthesis-of-peptides-unit/). The peptide moiety was synthesized on a microwave-assisted peptide synthesizer (Liberty Blue,CEM), using Rink amide Protide resin (0.56 mmol/g, CEM) as a solid support and a Fmoc/tBu strategy. Diisopropylcarbodiimide (DIC) and Oxyme were used as coupling reagents. After completion of the peptide moiety, Verteporfin (Medchem Express cat # HY-B0146) (2 equivalents) was manually introduced. CES-V was released from the solid support by treatment with TFA: CH_2_Cl_2_:TIS (95:2.5:2.5, v/v/v) for 1.5 h. The solvent was then evaporated under vacuum, and the peptide conjugates were precipitated with cold diethyl ether. The solid obtained was decanted and dissolved in a mixture of H_2_O:CH_3_CN (1:1, v/v) and lyophilized.

To generate the CES-V conjugate with a disulphide bridge between C1 and C9, a 1mM solution of the linear CES-V conjugate in H_2_O:CH_3_CN (1:1, v/v) was prepared and the pH was adjusted to 8 with a solution of 20% NH_4_Cl in H_2_O. The evolution of disulphide formation was monitored by HPLC and was completed after 12 h. CES-V was purified by semipreparative HPLC with a XBridge Peptide BEH C18 OBD Prep column (130 Å, 5 μm, 19 x100 mm), using H2O (1% CF_3_COOH) and CH_3_CN (1% CF_3_COOH) as eluents. Final pure CES-V was analyzed and characterized by HPLC and HPLC-MS.

### Animal models

AD animal experiments were conducted under an approved protocol of the Institutional Animal Care and Use Committee of Sanford Burnham Prebys Medical Discovery Institute. Nine-month-old transgenic C57BL/6 mice (models hAPP-J20) were used for peptide homing studies. Glioblastoma animal experiments were performed under approved protocols of the committee of Animal Experimentation of Estonian Ministry of Agriculture (Permit #159). Orthotopic glioblastoma (for peptide homing studies, receptor identification and anti-SNAP25 homing) were induced into female athymic nude mice by implanting 3 × 10^5^ U87MG cells in a volume of 2.5 µl intracranially in the right striatum of the brain (coordinates: 2 mm laterally and 2 mm posteriorly from bregma and at 2.5 mm depth). Tumors were allowed to develop for 9 days before performing an experiment.

For the angiogenic glioblastoma model, WT-GBM^38^, 7 × 10^5^ cells were stereotactically injected in the right striatum of Fox/Nu mice 2 mm lateral and 2 mm posterior of bregma at depth of 2.5 mm. The mice were used for homing studies 6-7 days after the injection.

For MCF10CA1a tumor induction, athymic nude mice were injected with 2× 10^6^ MCF10CA1a triple-negative breast tumor cells in 50 µL PBS in the mammary gland and the mice were used 14 days after the inoculation.

### Homing studies

For peptide homing, mice were intravenously injected with 30 nmoles of peptide dissolved in PBS and allowed to circulate for 60 min. Mice were then perfused intracardially with PBS and all major organs were isolated and fixed in 4% paraformaldehyde (PFA) at pH 7.4 overnight. The organs were then washed with PBS and placed in graded sucrose solutions overnight before optimal cutting temperature compound (OCT) embedding. Ten μm-thick sections were cut and analyzed by immunostaining. Slides were stained, without the Triton 0.2% permeabilization step, for FAM (rabbit anti-FAM, ThermoFisher, cat # A889), CD31 (BD Biosciences, cat # 553370), SNAP25 (LSBIO, cat # LS-B2392-50), PDGFRB (R&D, cat # AF1042), CD68 (Bio-Rad, cat # MCA1957GA), at 1/100 dilution overnight at +4C, and then stained at 1/200 using Alexa Fluor 647 goat anti-rabbit (Thermo Fisher Scientific, cat # A21245), Alexa Fluor 546 goat anti-rat antibody (Life Technologies, cat # A11081) or Alexa Fluor 546 donkey anti-goat (Thermofisher, cat # A-11056). Images were taken with a LSM710 Zeiss confocal.

### Affinity chromatography

To identify the receptor of the CES peptide, affinity chromatography was performed as described previously^9^ with slight modifications. Four intracranial U87MG tumors were excised, snap-frozen and crushed in liquid nitrogen using mortar and pestle. The powder was dissolved in lysis buffer (400 mM n-octyl-β-D-glucopyranoside, 2 mM MnCl_2_, 2 mM MgSO_4_, 2 mM CaCl_2_, 20× solution of EDTA-free Sigmafast Protease Inhibitor Cocktail Tablets (Sigma-Aldrich) in PBS) and the cleared lysate was incubated with streptavidin-coated magnetic beads (Dynabeads M-280 Streptavidin, Invitrogen) conjugated with CES peptide. The beads were washed with diluted lysis buffer (1:1 in PBS), followed by washes with 0.2 mM and 0.5 mM control peptide (Ac-CRKQGEAKC, disulfide cyclized) to remove non-specifically bound proteins. Finally, the bound proteins were eluted using 2 mM Ac-CES peptide. The eluted fractions and the fractions collected during wash steps were analyzed using mass spectrometry analysis at the institute of technology of the University of Tartu.

### Quartz crystal microbalance

The Quartz Crystal Microbalance (QCM) used was QCM200 system from Stanford Research Systems (Sunnyvale, CA, USA). The quartz crystals used have 5MHz resonant frequency and are deposited with a layer of Cr/Au (Cat # O100RX1, p/n 6-613, Stanford Research Systems). The crystal was mounted on the cell, washed with isopropanol, ethanol and mQ water. Then, it was incubated with 20mM Mercapto-propanesulfonate (MPS, Cat #251682, Sigma-Aldrich) in 10mM H_2_SO_4_ for 30 minutes, washed with mQ and then incubated with a 10µM (in monomer) of Poly (allylamine hydrochloride, Mw: 50000, PAH, Cat # 283223, Sigma-Aldrich) in mQ at pH 8 during 10 min and later washed with mQ. Then, human recombinant SNAP25 (Cat # ab74529, Abcam) or Bovine Serum Albumin (BSA, Cat # A7906, Sigma-Aldrich) were deposited at a concentration of 0.01mg/mL in mQ for 10 minutes, and then washed with mQ. Then, 500µL of PBS were placed in the cell and the baseline was recorded, the measurement was then paused and 50µL of a 300µM solution of CES or iRGD in PBS were gently deposited on the cell and the measurement was resumed. Then, the measurement was paused, the solution removed and placed 500µL of new PBS and the measurement resumed. The association and dissociation curves were fitted using TraceDrawer software (Ridgeview Instruments AB) using a one-to-one model, to obtain the association constant ka, the dissociation constant kd and the affinity constant K_D_.

### Molecular Dynamics

Receptor SNAP-25 was modeled using AlphaFold ^39^. CES peptide was built from its sequence: CYS GLU SER PRO LEU LEU SER GLU CYS NHE. Parameters used to describe molecular interactions of the SNAP25-CES complex were prepared with the module tLeap from the AmberTools package^40^. The amber forcefield ff14SB^41^ was used for the protein and peptides, and TIP3P model^42^ to describe explicit water molecules. K+ and Cl-ions were added to construct 0.15 M KCl solutions. Each system (receptor in water, peptides in water and complex in water) was enclosed in a cubic box, and periodic boundary conditions were used to perform the dynamics.

Simulation protocols consisted of 4000 steps of minimization with the steepest descent algorithm followed by 6000 steps of conjugate gradient algorithm. NVT ensemble was used and periodic boundary conditions were implemented with Particle Mesh Ewald (PME)^43^. A 10 Å cut-off was used. After minimization, the systems were heated to 298 K using Langevin dynamics^44^ with collision frequency of 2 ps^-1^. Shake algorithm^45^ was turned on. Isotropic position scaling (was used to get a pressure of 1 atm with a relaxation time of 1 ps. Systems were then equilibrated for 10 ns and a production run of 50 ns was performed for each one.

From molecular dynamics simulations a trajectory was generated, and cluster analysis was performed to determine the most representative conformations. Clustering was implemented using a distance metric (RMSD), and with the average-linkage algorithm.

Four cluster nodes were selected as representative structures for the receptor SNAP-25 and two cluster nodes for the CES peptide.

### Docking

The two representative structures for CES obtained from molecular dynamics were docked against the four representative structures obtained for SNAP-25 using a blind peptide-protein docking algorithm: HPEPDOCK^46^.

### MM-PBSA calculation

Relative binding free energies of the CES-SNAP-25 complex were calculated and compared against a control complex; the iRGD-SNAP25 complex. Molecular Mechanics-Poisson Boltzman Surface Area (MMPBSA) method^47^ was used to calculate the binding energies. Binding free energies for each complex were calculated for a 20 ns MD trajectory.

The SNAP25-Collagen VI complex was modelled using AlphaFold3^48^, yielding a predicted template modeling (pTM) score of 0.31 and an interface predicted template modeling (ipTM) score of 0.15. While these scores are low, they reflect the accuracy of the entire structure. In this case, three regions, including the binding site, are predicted with high accuracy, while the rest of the protein, distant from the binding site, is predicted with lower accuracy, thus reducing the overall score.

### *In vitro* photodynamic therapy

30.000 cells were seeded on 96 well plates and left overnight to attach. Then, the cells were incubated with the conjugates at 3uM (by adding 10µL of a 30µM solution), for 30 min at room temperature. Then, the wells were washed 3 times with full media, placed in new media and irradiated with a dose of 10 Joules using a 160 mW/cm^2^ irradiance, using a 690nm laser. Then, the cells were left in culture for additional 24 hours and later the viability was assayed using 20µL of MTS/PMS (Promega, Catalog# MTS: G109A PMS: 611OA) incubating 2 hours at +37 °C in a humidified atmosphere containing 5% CO_2_ and reading absorbance at 492 nm.

### Flow cytometry

The cells were harvested mechanically by adding 5mL of PBS and using a cell scraper to detach the cells. Once cells were fully detached, they were resuspended with a pipette, transferred to a collection tube, and centrifuged for 6 minutes at 250G. Upon centrifugation, the supernatant was removed and 200µL of Blocking buffer (PBS with 5% BSA (W/V) + 5% FBS(V/V)) were added for every 200,000 cells. The cells were resuspended in blocking buffer and left on agitation at 4°C for 30 minutes. Then, 2µL of the antibody (FITC mouse anti-SNAP25, Biolegend, cat # 836313) or its isotype control (FITC-mouse IgG isotype control, Biolegend, cat# 400132) were added, the tubes were protected with aluminum foil and incubated at 4°C for 30 minutes on agitation. Then, the samples were washed twice by centrifuging for 6 minutes at 250G, discarding the supernatant and resuspending in PBS. Finally, the cells were brought to a total volume of 200µL in PBS and read in a Guava easyCyte flow cytometer.

### *In vitro* immunofluorescence microscopy

U87MG cells were cultured on µ-slide 8 well high ibiTreat (Ibidi, Cat # 80806) slides for 48 hours on 200 µL of culture media, after which the media was removed, and one wash was performed using PBS. The cells were then fixed by using 4% paraformaldehyde for 10 minutes at room temperature. Once fixed, cells were washed four times with PBS and incubated with blocking buffer (PBS with 5% FBS (V/V) and 5% BSA (W/V)) for 1 h at room temperature. After, blocking buffer was removed and cells were incubated with A488 anti-SNAP25 (Biolegend, Cat # 836313) diluted 1/300, in diluted blocking buffer (1/5 in PBS) or its isotype control (Biolegend, Cat # 400132) at 4°C overnight. Next, cells were washed 4 times with PBS and incubated with DAPI (Invitrogen, Cat # 10184322) 5μg/mL in PBS for 5 minutes at room temperature. Finally, 4 washes were performed using PBS and the slide was imaged in PBS, in a confocal microscope (ZEISS LSM780).

### Binding of SNAP25 to ECM array

The ECM Select® Array Kit slide (Cellsystems Cat # 5170-1EA) was assembled onto the slide holder. The slide was washed three times with PBS, leaving 4 minutes intervals between each wash, followed by two washes with PBST (Tween-20 0.05% V/V in PBS). The slide was then incubated during one hour at room temperature with blocking buffer (5% W/V BSA in PBST+ 5%V/V FBS). After, it was incubated overnight at 4°C with SNAP25 human recombinant protein (Abcam Cat # 74529) diluted 1/200 in diluted blocking buffer (1/5 in PBST). Subsequently, the slide was washed three times with PBST and incubated with A488 anti-SNAP25 (Biolegend, Cat # 836313), diluted 1/200 in diluted blocking buffer, overnight at 4°C. Finally, the slide was washed three times with PBST leaving 4 minutes intervals, followed by 3 washes with PBS. The slide was imaged using a fluorescence microscope (EVOS™ M7000) in the green channel.

### Surface Plasmon Resonance

SPR analyses were conducted at 20 °C in an OpenSPRTM instrument (Nicoya), using a high sensitivity NTA sensor chip (Nicoya Cat# NICSEN-HS-8-NTA). Surface conditioning was performed on channels 1 and 2 with 200µL of 10mM HCl injected at a flow rate of 150µl per minute. Followed by imidazole 200mM at 20µl per minute, and EDTA 350Mm at 100µl per minute. Once conditioned, the sensor surface was activated with NiCl2 40mM injected at 20µl per minute. SNAP25 human recombinant protein (Abcam Cat# 74529) at a concentration of 150µg/mL was immobilized using a flow rate of 20µl per minute only through channel 2. Capture analysis was performed injecting Collagen 20µg/ml (collagen 50µg/ml in 500mM of acetic acid diluted to 20µg/ml in PBS) through both channels, with flow rate of 20µl per minute. Between each injection, 1ml of PBS was flushed though the loop. Finally, the device was washed with 200µL of PBS at 20µl per minute though both channels. As a binding control, recombinant mouse macrophage colony-stimulating factor (M-CSF) (28 kDa, with 9-His tag) (Biolegend, Cat# 576406) at 10µg/mL was immobilized using a flow rate of 20µl per minute only though channel 2, and capture analysis was performed using the same settings as described previously. The data was fitted with TraceDrawer using a one-to-one model.

### Cell culture on collagens V and VI

Bovine Type VI Collagen-Solution (Southern Biotech Cat # 1300-02S) was diluted to 50µg/ml in 500mM of acetic acid. Human type V Collagen-Lyophilized (Southern Biotech, Cat # 1270-01) was reconstituted to 50µg/ml in 500mM of acetic acid. To coat the 96-well plates 5µg/cm^2^ of 50µg/ml collagen VI or V was added and incubated at room temperature for 1 h. After which collagen was removed and the plate washed twice with PBS.

To evaluate cell adhesion of U87MG cells on collagen VI, the cells were seeded and cultured in 100µL in the presence or absence of mouse anti-SNAP25 antibody (Biolegend, Cat # 836303) at a concentration of 5µg/mL or 30µM of CES. Then, 24 hours later, media was changed, and cells were cultured for additional 48 hours. Then, cells were fixed with 4% paraformaldehyde at room temperature for 10 minutes, followed by incubation with blocking buffer (PBS with 5% FBS (V/V) and BSA (W/V)) for 1 h at room temperature. Once blocking was completed, cells were washed three times with PBS and incubated with DAPI (Invitrogen, Cat # 10184322) 5μg/mL in PBS for 5 minutes at room temperature. Finally, 4 washes were performed using PBS and the cells were imaged using a fluorescence microscope (EVOSTM M7000). Cell density was calculated by counting the nuclei in the images (using ImageJ or manually). The resulting cell count was divided by the area of the images (1.2mm^2^).

### Anti-SNAP25 homing

Glioblastoma and healthy mice were injected intraperitoneally with rabbit polyclonal anti-human/mouse SNAP25 (Thermofisher, cat # PA5-19701), (33µg in 500µL of PBS) and circulated for 24 hours. Then, mice were perfused with 20mL of PBS and the tissues collected and placed in 4% PFA overnight, then washed during 1 h at room temperature in PBS to remove PFA and placed in gradients of sucrose (first 15% then 30%, each overnight at +4). The tissues were then placed in OCT, sectioned and stained without permeabilization, using A546 goat anti-rabbit (Thermofisher, Catalog # A-11010) (to visualize the administered anti-SNAP), and A647 goat anti-mouse (Thermofisher, Catalog # A-21235) (to visualize endogenous mouse IgG), both at dilution 1/200.

## Notes

### Competing Interest Statement

ER, AM and PS are co-inventors on patents describing the reported peptides.

## References

1. Ruoslahti, E. Molecular ZIP codes in targeted drug delivery. Proceedings of the National Academy of Sciences 119, e2200183119 (2022).

2. Fraguas-Sánchez, A. I., Lozza, I. & Torres-Suárez, A. I. Actively Targeted Nanomedicines in Breast Cancer: From Pre-Clinal Investigation to Clinic. Cancers (Basel) 14, 1198 (2022).

3. Schliemann, C. et al. Three clinical-stage tumor targeting antibodies reveal differential expression of oncofetal fibronectin and tenascin-C isoforms in human lymphoma. Leukemia Research 33, 1718–1722 (2009).

4. Sugahara, K. N. et al. Coadministration of a tumor-penetrating peptide enhances the efficacy of cancer drugs. Science 328, 1031–1035 (2010).

5. Dean, A. et al. Dual αV-integrin and neuropilin-1 targeting peptide CEND-1 plus nab-paclitaxel and gemcitabine for the treatment of metastatic pancreatic ductal adenocarcinoma: a first-in-human, open-label, multicentre, phase 1 study. The Lancet Gastroenterology & Hepatology 7, 943–951 (2022).

6. Scodeller, P. & Asciutto, E. K. Targeting Tumors Using Peptides. Molecules 25, 808 (2020).

7. Bessone, M. I. D. et al. IRGD-guided tamoxifen polymersomes inhibit estrogen receptor transcriptional activity and decrease the number of breast cancer cells with self-renewing capacity. Journal of Nanobiotechnology 17, (2019).

8. Mann, A. P. et al. A peptide for targeted, systemic delivery of imaging and therapeutic compounds into acute brain injuries. Nat Commun 7, 11980 (2016).

9. Mann, A. P. et al. Identification of a peptide recognizing cerebrovascular changes in mouse models of Alzheimer’s disease. Nature Communications 8, (2017).

10. Pleiko, K. et al. In vivo phage display: identification of organ-specific peptides using deep sequencing and differential profiling across tissues. Nucleic Acids Res 49, e38 (2021).

11. Blouw, B. et al. The hypoxic response of tumors is dependent on their microenvironment. Cancer Cell 4, 133–146 (2003).

12. Du, R. et al. Matrix metalloproteinase-2 regulates vascular patterning and growth affecting tumor cell survival and invasion in GBM. Neuro-Oncology 10, 254–264 (2008).

13. Gavish, A. et al. Hallmarks of transcriptional intratumour heterogeneity across a thousand tumours. Nature 618, 598–606 (2023).

14. Yu, C. et al. Association between SNAP25 and human glioblastoma multiform: a comprehensive bioinformatic analysis. Biosci Rep 40, BSR20200516 (2020).

15. Huang, Q. et al. SNAP25 Inhibits Glioma Progression by Regulating Synapse Plasticity via GLS-Mediated Glutaminolysis. Frontiers in Oncology 11, (2021).

16. Baltus, R. E., Carmon, K. S. & Luck, L. A. Quartz Crystal Microbalance (QCM) with Immobilized Protein Receptors: Comparison of Response to Ligand Binding for Direct Protein Immobilization and Protein Attachment via Disulfide Linker. Langmuir 23, 3880–3885 (2007).

17. Naklua, W., Suedee, R. & Lieberzeit, P. A. Dopaminergic receptor–ligand binding assays based on molecularly imprinted polymers on quartz crystal microbalance sensors. Biosensors and Bioelectronics 81, 117–124 (2016).

18. Janshoff, A. & Steinem, C. Label-free detection of protein-ligand interactions by the quartz crystal microbalance. Methods Mol Biol 305, 47–64 (2005).

19. Migoń, D., Wasilewski, T. & Suchy, D. Application of QCM in Peptide and Protein-Based Drug Product Development. Molecules 25, 3950 (2020).

20. Dunér, G. et al. Signal enhancement in ligand–receptor interactions using dynamic polymers at quartz crystal microbalance sensors. Analyst 141, 3993–3996 (2016).

21. Flexer, V., Forzani, E. S., Calvo, E. J., Ludueña, S. J. & Pietrasanta, L. I. Structure and Thickness Dependence of “Molecular Wiring” in Nanostructured Enzyme Multilayers. Anal. Chem. 78, 399– 407 (2006).

22. Scodeller, P. et al. Layer-by-Layer Self-Assembled Osmium Polymer-Mediated Laccase Oxygen Cathodes for Biofuel Cells: The Role of Hydrogen Peroxide. J. Am. Chem. Soc. 132, 11132– 11140 (2010).

23. Scodeller, P. et al. Wired-Enzyme Core−Shell Au Nanoparicle Biosensor. J. Am. Chem. Soc. 130, 12690–12697 (2008).

24. Sarvas, H. O., Seppälä, I. J. T., Tähtinen, T., Péterfy, F. & Mäkelä, O. Mouse IgG antibodies have subclass associated affinity differences. Molecular Immunology 20, 239–246 (1983).

25. Hagiwara, N., Kadono, N., Miyazaki, T., Maekubo, K. & Hirai, Y. Extracellular syntaxin4 triggers the differentiation program in teratocarcinoma F9 cells that impacts cell adhesion properties. Cell Tissue Res 354, 581–591 (2013).

26. Shirai, K. et al. Extracellularly Extruded Syntaxin-4 Binds to Laminin and Syndecan-1 to Regulate Mammary Epithelial Morphogenesis. Journal of Cellular Biochemistry 118, 686–698 (2017).

27. Holz, R. W. & Bittner, M. A. Roles for the SNAP25 linker domain in the fusion pore and a dynamic plasma membrane SNARE “acceptor” complex. J Gen Physiol 152, e202012619 (2020).

28. Gonzalo, S., Greentree, W. K. & Linder, M. E. SNAP-25 Is Targeted to the Plasma Membrane through a Novel Membrane-binding Domain *. Journal of Biological Chemistry 274, 21313–21318 (1999).

29. Zhu, Q., Yamakuchi, M. & Lowenstein, C. J. SNAP23 Regulates Endothelial Exocytosis of von Willebrand Factor. PLoS One 10, e0118737 (2015).

30. Fogal, V., Sugahara, K. N., Ruoslahti, E. & Christian, S. Cell surface nucleolin antagonist causes endothelial cell apoptosis and normalization of tumor vasculature. Angiogenesis 12, 91– 100 (2009).

31. Fogal, V., Zhang, L., Krajewski, S. & Ruoslahti, E. Mitochondrial/Cell-Surface Protein p32/gC1qR as a Molecular Target in Tumor Cells and Tumor Stroma. Cancer Research 68, 7210– 7218 (2008).

32. Säälik, P. et al. Peptide-guided nanoparticles for glioblastoma targeting. Journal of Controlled Release 308, 109–118 (2019).

33. Silverman, J. M. et al. CNS-derived extracellular vesicles from superoxide dismutase 1 (SOD1)G93A ALS mice originate from astrocytes and neurons and carry misfolded SOD1. J Biol Chem 294, 3744–3759 (2019).

34. Matsuguchi, S. & Hirai, Y. Syntaxin4, P-cadherin, and CCAAT enhancer binding protein β as signaling elements in the novel differentiation pathway for cultured embryonic stem cells. Biochemical and Biophysical Research Communications 672, 27–35 (2023).

35. Turtoi, A. et al. Accessibilome of Human Glioblastoma: Collagen-VI-alpha-1 Is a New Target and a Marker of Poor Outcome. J. Proteome Res. 13, 5660–5669 (2014).

36. Paulus, W., Roggendorf, W. & Schuppan, D. Immunohistochemical investigation of collagen subtypes in human glioblastomas. Vichows Archiv A Pathol Anat 413, 325–332 (1988).

37. Cha, J. et al. Glioma Cells Secrete Collagen VI to Facilitate Invasion. bioRxiv 2023.12.12.571198 (2023) doi:10.1101/2023.12.12.571198.

38. Blouw, B. et al. The hypoxic response of tumors is dependent on their microenvironment. Cancer Cell 4, 133–146 (2003).

39. Varadi, M. et al. AlphaFold Protein Structure Database: massively expanding the structural coverage of protein-sequence space with high-accuracy models. Nucleic Acids Research 50, D439– D444 (2022).

40. Case, D. A. et al. AmberTools. J. Chem. Inf. Model. 63, 6183–6191 (2023).

41. Maier, J. A. et al. ff14SB: Improving the Accuracy of Protein Side Chain and Backbone Parameters from ff99SB. J. Chem. Theory Comput. 11, 3696–3713 (2015).

42. Jorgensen, W. L., Chandrasekhar, J., Madura, J. D., Impey, R. W. & Klein, M. L. Comparison of simple potential functions for simulating liquid water. The Journal of Chemical Physics 79, 926– 935 (1983).

43. Darden, T., York, D. & Pedersen, L. Particle mesh Ewald: An N⋅log(N) method for Ewald sums in large systems. The Journal of Chemical Physics 98, 10089–10092 (1993).

44. Paterlini, M. G. & Ferguson, D. M. Constant temperature simulations using the Langevin equation with velocity Verlet integration. Chemical Physics 236, 243–252 (1998).

45. Ryckaert, J.-P., Ciccotti, G. & Berendsen, H. J. C. Numerical integration of the cartesian equations of motion of a system with constraints: molecular dynamics of n-alkanes. Journal of Computational Physics 23, 327–341 (1977).

46. Zhou, P., Jin, B., Li, H. & Huang, S.-Y. HPEPDOCK: a web server for blind peptide–protein docking based on a hierarchical algorithm. Nucleic Acids Res 46, W443–W450 (2018).

47. Miller, B. R. I. et al. MMPBSA.py: An Efficient Program for End-State Free Energy Calculations. J. Chem. Theory Comput. 8, 3314–3321 (2012).

48. Abramson, J. et al. Accurate structure prediction of biomolecular interactions with AlphaFold 3. Nature 630, 493–500 (2024).

