## Supporting information for "A CYCLIC PEPTIDE TARGETS GLIOBLASTOMA BY BINDING TO ABERRANTLY EXPOSED SNAP25"

**TABLE 1:**
**Proteins identified by CES peptide-affinity chromatography and mass spectrometry (MS) analysis**

| Protein ID | Protein name | Gene name |
|---|---|---|
| Q3UV17 | Keratin, type II cytoskeletal 2 oral | Krt76 |
| P62806 | Histone H4 | Hist1h4a |
| Q02257 | Junction plakoglobin | Jup |
| G3UW94 | MCG14259, isoform CRA_b | U2af1 |
| P52479 | Ubiquitin carboxyl-terminal hydrolase 10 | Usp10 |
| Q9JHW9 | Aldehyde dehydrogenase family 1 member A3 | Aldh1a3 |
| A0A0G2JG70 | Zinc finger and SCAN domain-containing 25 | Zscan25 |
| Q05816 | Fatty acid-binding protein, epidermal | Fabp5 |
| P10107 | Annexin A1 | Anxa1 |
| A2AFI4 | RNA-binding motif protein, X chromosome (Fragment) | Rbmx |
| F6X2U6 | Serine/arginine-rich-splicing factor 11 (Fragment) | Srsf11 |
| Q5SXR6 | Clathrin heavy chain | Cltc |
| A0A1Y7VKP8 | NADH dehydrogenase [ubiquinone] iron-sulfur protein 6, mitochondrial | Ndufs6 |

| | | |
|---|---|---|
| Q9QYG0-2 | Isoform 2 of Protein NDRG2 | Ndrg2 |
| Q8VIJ6 | Splicing factor, proline- and glutamine-rich | Sfpq |
| P63260 | Actin, cytoplasmic 2 | Actg1 |
| Q6NXW6 | Cell cycle checkpoint protein RAD17 | Rad17 |
| P17897 | Lysozyme C-1 | Lyz1 |
| P61982 | 14-3-3 protein gamma | Ywhag |
| P60879-2 | Isoform 2 of Synaptosomal-associated protein 25 | Snap25 |

**Table S1. Proteomics hits ordered by the ratio of CES-eluted fraction/Ctrl peptide eluted fraction**

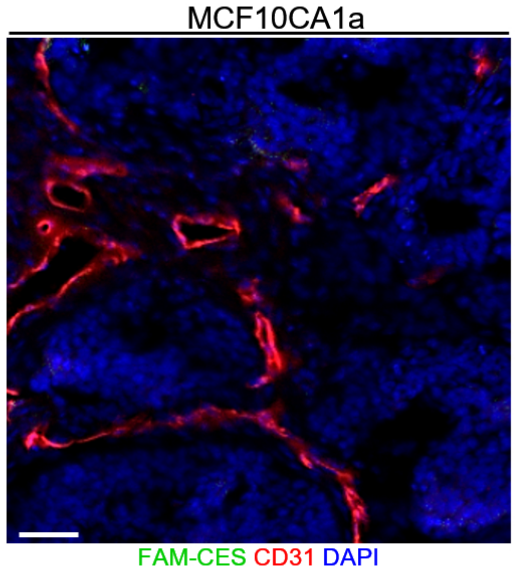


**Figure S1.** FAM-CES does not home to MCF10CA1a breast cancer. Homing of intravenously administered FAM-CES in mice bearing orthotopic MCF10CA1a tumors (C) (n=3). 30 nmoles of FAM-CES were administered intravenously and allowed to circulate for 60 min. The mice were perfused, and the tumors were fixed, sectioned, and stained for FAM (green) and CD31. Scale bar= 50µm.

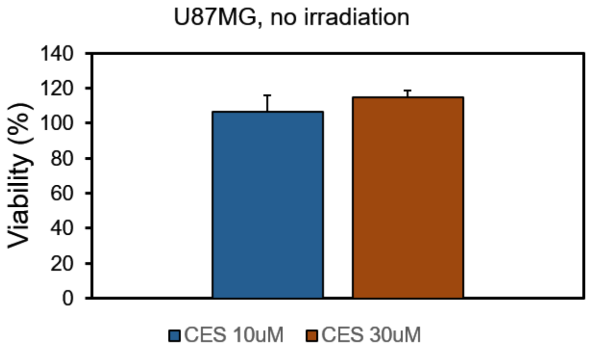


**Figure S2. In vitro viability with CES in U87MG cells in absence of laser irradiation.** Cells were seeded on 96-well plates and 24 hours later incubated with CES for 30 min at 37°C, followed by three washes with media, the cells were cultured for additional 24 h and the viability was assessed using MTS assay.

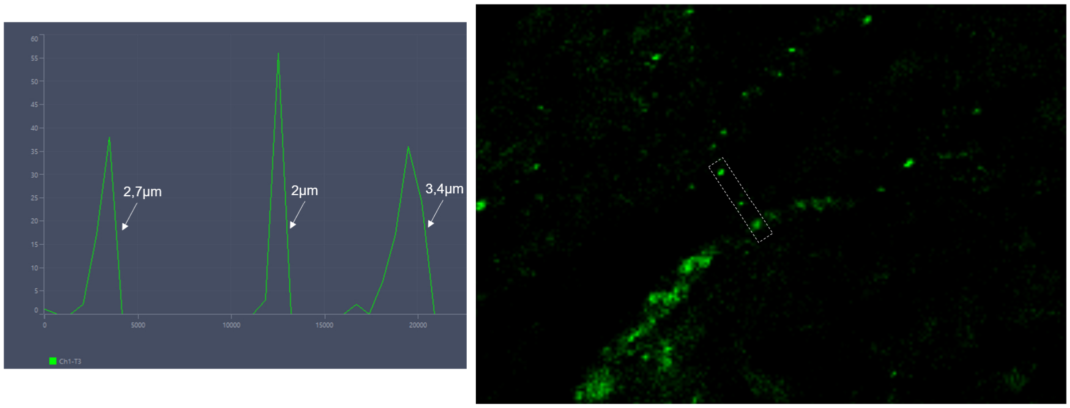


**Figure S3**. Size of particulate SNAP25 staining (green) observed on CD31⁺ blood vessels in U87MG tumors.

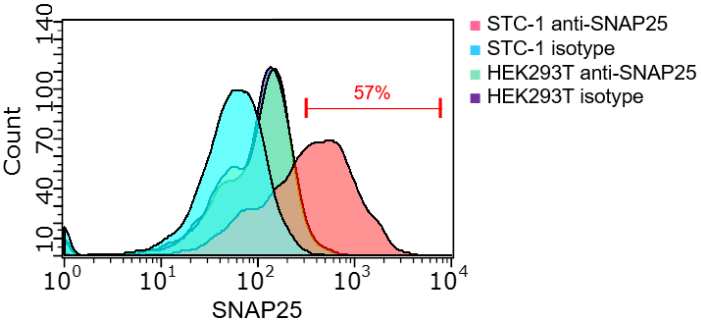


**Figure S4.** Expression of superficial SNAP25 in STC-1 and HEK293T cells using live cell cytometry performed at +4C and in the absence of any detergent.

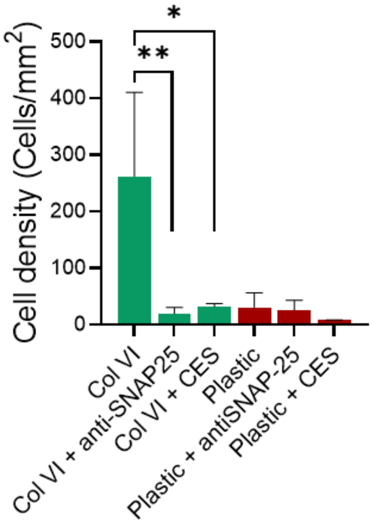


**Figure S5.** Cellular attachment of U87MG cells on collagens VI. Cells were seeded on collagen VI or plastic, in the presence or absence of anti-SNAP25 (5µg/mL) or CES (30µM) for 24h. Then, cells were washed and continued to grow for an additional 48h. Cells were then washed, fixed, and stained for DAPI and the number of cells in an area of 1,2mm$^2$ were counted with a fluorescent microscope. Bars represent mean + SE.
